# I know that I don’t know: Structural and functional connectivity underlying meta-ignorance in pre-schoolers

**DOI:** 10.1101/450346

**Authors:** Elisa Filevich, Caroline Garcia Forlim, Carmen Fehrman, Carina Forster, Markus Paulus, Yee Lee Shing, Simone Kühn

**Author notes:** Yee Lee Shing and Simone Kühn should be considered joint senior author.

## Abstract

**Research Highlights:** [1] Children develop the ability to report that they do not know something at around five years of age.

[2] Children who could correctly report their own ignorance in a partial-knowledge task showed thicker cortices within medial orbitofrontal cortex.

[3] This region was functionally connected to parts of the default-mode network.

[4] The default-mode network might support the development of correct metacognitive monitoring.

**Abstract:** Metacognition plays a pivotal role in human development. The ability to realize that we do not know something, or meta-ignorance, emerges after approximately five years of age. We aimed at identifying the brain systems that underlie the developmental emergence of this ability in a preschool sample.

Twenty-four children aged between five and six years answered questions under three conditions of a meta-ignorance task twice. In the critical *partial knowledge* condition, an experimenter first showed two toys to a child, then announced that she would place one of them in a box behind a screen, out of sight from the child. The experimenter then asked the child whether or not she knew which toy was in the box.

Children who answered correctly both times to the metacognitive question in the partial knowledge condition (n=9) showed greater cortical thickness in a cluster within left medial orbitofrontal cortex than children who did not (n=15). Further, seed-based functional connectivity analyses of the brain during resting state revealed that this region is functionally connected to the medial orbitofrontal gyrus, posterior cingulate gyrus and precuneus, and mid- and inferior temporal gyri.

This finding suggests that the default mode network, critically through its prefrontal regions, supports introspective processing. It leads to the emergence of metacognitive monitoring allowing children to explicitly report their own ignorance.

## Introduction

Metacognition, or the ability to monitor one’s own mental states and processes, is a crucial cognitive function that enables flexible and adaptive behaviour. It has been shown to be a strong predictor of cognitive development and, in particular, school achievements (Schneider, 2008; Williams et al., 2002). For example, by determining what we know, or do not know, we can decide whether to act quickly or seek more information before committing to a course of action (Desender, Boldt, & Yeung, 2018; Yeung & Summerfield, 2012). “Metacognition” is a broad term, and may be measured through a wide variety of behavioural paradigms. These include, for example, the monitoring and explicit report of one’s own memory (Chua, Pergolizzi, & Weintraub, 2014), perception (Fleming, Weil, Nagy, Dolan, & Rees, 2010), or focus of attention (Whitmarsh, Barendregt, Schoffelen, & Jensen, 2014; Whitmarsh, Oostenveld, Almeida, & Lundqvist, 2017), as well as the implicit control of attentional resources (Kentridge & Heywood, 2000), error monitoring (Charles, Opstal, Marti, & Dehaene, 2013) or allocation of study time (Son & Metcalfe, 2000).

Partially different regions within the prefrontal cortex (PFC) have been found to support different aspects of metacognitive monitoring (Dehaene, Lau, & Kouider, 2017; Fleming & Dolan, 2012). Yet, however, from a developmental point of view, two core questions remain unanswered. First, what is the specific role of these frontal regions and how do they relate to other structures? Second, how does the brain’s monitoring ability develop? In the past few years, brain imaging experiments on adult volunteers (e.g. Baird, Smallwood, Gorgolewski, & Margulies, 2013; Fleming et al., 2010; McCurdy et al., 2013) aimed at answering the first question, whereas behavioural experiments in developing populations aimed at answering the second (e.g. Balcomb & Gerken, 2008; Goupil, Romand-Monnier, & Kouider, 2016; S. Kim, Paulus, Sodian, & Proust, 2016; Rohwer, Kloo, & Perner, 2012; Vo, Li, Kornell, Pouget, & Cantlon, 2014). As a result, these two questions have been largely studied independently of one another. Here, we aimed at bridging these two research approaches to characterize the development of the neural correlates of the emergence metacognitive monitoring in early childhood.

To the best of our knowledge, only one recent study investigated the neural bases of metacognition in a developmental sample. Fandakova et al. (2017) related longitudinal changes in cortical structure with changes in meta-memory ability in the transition from late childhood into adolescence (7- to 12-year-old children). They found that the rate of cortical thinning in anterior insula and of cortical thickening in ventromedial PFC were related to changes in meta-memory monitoring ability over time. This result suggests that, as in adults, the PFC plays a crucial role in metacognitive monitoring ability already during childhood. While this study provides unique evidence on the neural correlates related to metacognitive development in late childhood and adolescents, it remains unclear, which neural networks support the emergence of metacognitive abilities in early childhood. Our study attempts to address this question.

In what follows, we first briefly review existing behavioural results on the emergence of metacognition before turning to the task that we employed in this study.

### When do metacognitive abilities develop?

Different aspects of metacognitive monitoring have been shown to develop at different ages. A particularly useful distinction is between implicit metacognition —measured through its effect on behaviour, potentially present from early on— and explicit metacognition, measured through the accuracy of explicit verbal judgements, emerging in the preschool years (Proust, 2013). For example, infants persist in their answers for a longer time after correct than after incorrect choices by 12 months and can regulate their waiting times for a reward in a manner that corresponds to their probability of being correct by 18 months (Goupil & Kouider, 2016). Moreover, Kim et al. (2016) showed that 3- and 4-year olds are able to recognize that they do not have a piece of information by choosing not to inform a third person about it. Crucially however, when Kim et al. explicitly asked the same children whether they themselves had this piece of information, children often (incorrectly) said that they did. It has been suggested that these processes are based on data-driven cues during the learning or performance processes itself (Koriat, Nussinson, Bless, & Shaked, 2008; Proust, 2013).

Explicit metacognition, on the other hand, refers to our ability to reflect on our cognitive processes and state our (lack of) knowledge. For example, Socrates famous sentence “I know that I do not know” is a prototypical explicit metacognitive statement. Classical research has shown that young children tend to equate knowing with seeing for both others and themselves, and develop the ability to distinguish between the two concepts after 4 years (Pratt & Bryant, 1990; Taylor, 1988; Wimmer, Hogrefe, & Perner, 1988). More complex metacognitive knowledge shows a more protracted development (Weil et al., 2013). These metacognitive abilities have been shown to predict school achievements (Schneider & Pressley, 1997). Thus, explicit metacognitive knowledge seems to emerge in the late preschool years.

### How do metacognitive abilities develop?

Two main hypotheses have been put forward to explain the emergence of (explicit) metacognitive ability. One notion (the “*Simulation theory*”) is that children learn to recognize their *own* mental states by building on the neural systems that monitor the mental states of *others* (i.e., theory of mind (ToM) e.g (Goldman, 2006; Harris, 1992). Some findings suggested indeed positive relations between early ToM competencies and later metacognitive abilities (Lecce, Demicheli, Zocchi, & Palladino, 2015; Lockl & Schneider, 2007). A different notion, stemming from the so called *theory-theory* suggests instead that children rely on lay theories and rules that are applied to self and other in order to understand mental states (e.g. Gopnik & Wellman, 1994; Perner, 1991) and that, in the way of Bayesian observers, children inform and narrow their priors as they learn to understand the world (Gopnik & Wellman, 2012). More concretely, in the case of ToM, these theories consist of general principles that can explain behaviour, e.g. “people act to satisfy their desires according to their beliefs” (Apperly, 2008).

To understand how metacognitive ability develops in the context of these two main contrasting theories, Rohwer et al (2012) designed a meta-ignorance task in which children had to evaluate what they knew: an experimenter placed a toy inside a box either in plain sight or out-of-sight from a child and asked her whether she knew, or did not know, which toy was in the box. Rohwer et al. asked this question in three conditions that differed in terms of the epistemic state of the child. Two of these conditions posed no serious challenge for children as young as 2-3 years old. Children this age could answer correctly in situations in which they had either full informational access, or none at all. However, in the key *partial knowledge* condition children had seen two possible toys but did not see which of them the experimenter had placed in a box. Crucially, in this condition children cannot provide a correct answer based on a simple seeing-is-knowing rule (cf. Wimmer, Hogrefe, & Perner, 1988). Here, it was found that children under 6 years old had great difficulty recognizing their own ignorance. In contrast, children aged 5 *can* correctly judge the mental states of others in situations of partial informational access (Pillow, Hill, Boyce, & Stein, 2000; Ruffman, 1996). Therefore, Rohwer et al. argued that children follow cues to answer metacognitive questions concerning their *own* knowledge: it will suffice that they can produce *a* plausible answer to a question (regardless of its accuracy) for children to judge that they know the answer to the question, regardless of whether this answer is correct. It has been suggested that this indicates the emergence of a mature understanding of knowledge by around 6 years (Kloo, Rohwer, & Perner, 2017).

In order to clarify the neurocognitive mechanisms that subserve the emergence of meta-ignorance in early childhood we sought to identify the brain regions supporting it. Given the role of the PFC for metacognitive monitoring in late childhood and adolescents (Fandakova et al., 2017), we hypothesized a relationship between cortical thickness in PFC and metacognitive ability. Moreover, to contribute to the developmental debate on the relations between ToM and metacognition, we then compared our results with the brain mechanisms that have been related to ToM (for a review see Schurz, Radua, Aichhorn, Richlan, & Perner, 2014).

## Methods

### Participants

For this study we tested children who were participating in the first wave of an ongoing longitudinal study (Hippokid; Brod, Bunge, & Shing, 2017). Twenty-four 5- and 6-year olds (mean age (±*SD*): 5.49 ± 0.4, 14 boys) were included. Children did a meta-ignorance task and a cognitive battery (see below). We tested five additional children but excluded them from analyses due to missing data in one or more of the tasks from the cognitive battery: One child did not do the working memory task, one did not do the cognitive control task, two did not understand or complete the reasoning task, and one did not answer to both repetitions of the partial knowledge.

The HippoKID study aimed at studying the effect of schooling on cognitive development and followed five-year old pre-schoolers longitudinally over two years. Children were recruited through advertisements in kindergartens, newspapers, and Internet forums for parents. Participating children received an honorarium of €10 per hour. All were native German speakers, had no history of psychiatric or neurological disorders or developmental delays (based on parental report), and were born with more than 37 weeks of gestational age. Most children belonged to families with high socioeconomic status.

The testing session lasted approximately 90 minutes, included cognitive testing and approximately 20 minutes of magnetic resonance imaging (MRI). To prevent any possible anxiety and excessive movement during MR image acquisition, we let children get accustomed to the scanner by spending time inside a mock scanner that looked and sounded exactly like the real one. Further, an experimenter stood next to the children while they lay on the scanner.

The German Psychological Society (DGPs) approved the study and the children’s parents or legal guardians gave written informed consent. Procedures conformed to the Declaration of Helsinki.

### Behavioural tasks

### Meta-ignorance task

We operationalized explicit metacognition following closely a paradigm developed by Rohwer et al. (2012). The task included three epistemic conditions that differed in terms of how much knowledge a child had about a toy hidden in a box (see figure 1.A). In all three conditions, the experimenter, sitting across a table from a child, put one of two toys inside a cardboard box, and then asked the child a series of questions about her knowledge. In the *complete-knowledge* condition, the experimenter first showed two toys to the child and asked her to name them. If she could name them correctly, the experimenter announced that she would place one of the toys in a box with a lid (29 × 18 × 11.5 cm), and did so in plain sight of the child. She then asked the child: “Do you know which toy is in the box, or do you not know?” (In German: “*Weißt du, welches Spielzeug in dem Karton ist, oder weißt du es nicht?*”). We call this the *knowledge* question. Depending on the answer, the experimenter asked follow-up questions. If the child said that she knew which toy was in the box, the experimenter asked, first: “O.K., then tell me which toy is in the box”, then the *confirmation* question: “Do you really know, or are you guessing?” (“*Weißt du das wirklich, oder rätst nur?*”); and finally, “How do you know that the [toy’s name] is in the box?” If, instead, the child said that she did not know which toy was in the box, the experimenter would ask: “Why don’t you know which toy is inside the box?” The procedure for the other two conditions was identical save for what the experimenter showed to the child. In the *no-knowledge* condition, the experimenter did not show the two toys to the child, before announcing that she would place one of them inside the box, behind the partition screen. Hence, in the *no-knowledge* condition, the child had seen neither of the toys. In the crucial *partial-knowledge* condition, the experimenter showed the two toys to the child and asked her to name them, but before placing one of the toys in the box, the experimenter put a black partition screen (60 × 39 cm) that occluded the child’s view of the toys and box. Hence, in the *partial-knowledge* condition, the child knew what two toys the experimenter could have put in the box, but did not know which one.

All children completed the three tasks twice in a fixed order: *complete-, no-, partial-, partial-, no-, complete-knowledge*. For each child, the experimenter randomly drew one of four predetermined sequences of toys and followed it. Eight different toys (see figure 1.B) were available for the two repetitions of the two different conditions (*complete-* and *partial-knowledge*) that required two toys each. One child could not name one of the toys, so the experimenter replaced it with an additional toy available.

We recorded a video of the testing sessions for all but three children, due to technical problems. We coded the responses to each of the questions as correct or incorrect.

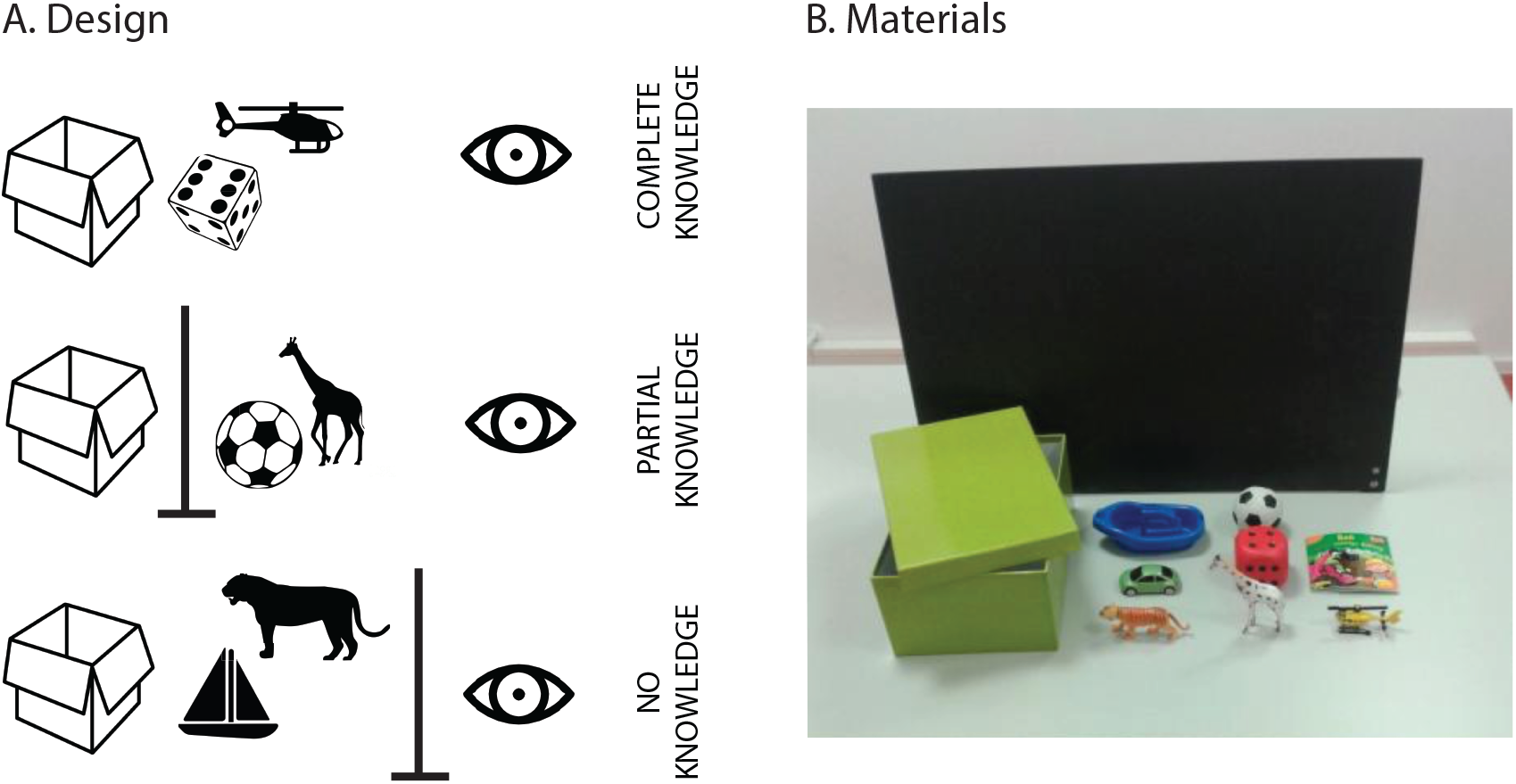

## Cognitive control - Hearts and flowers task

We operationalized cognitive control using the “hearts and flowers” task (Davidson, Amso, Anderson, & Diamond, 2006). The details have been described elsewhere (Brod et al., 2017). Briefly, the task included three conditions with 20 trials each. On each trial of the congruent condition (always presented during the first block of the task), an image of a heart appeared either on the right or left side of a computer screen and children had to press a key on the same side as the heart was displayed (with the corresponding right or left hand). The image was displayed for 1.5 s, and the trial ended 2 s after image onset. In the incongruent condition (always presented during the second block) an image of a flower appeared on either side of the screen, and children had to press a key on the side opposite to the flower. In the mixed condition (always presented in the third block), heart- and flower-stimuli were interleaved and children had to press a keyboard key on the same side of a heart displayed but on the opposite side of a flower displayed. This task requires sustained attention, maintenance of task rules in working memory and, in the mixed condition, inhibitory control and cognitive flexibility. We calculated each child’s accuracy (as the rate of correct responses) in the mixed condition, as it has been suggested that accuracy is a more sensitive measure of performance for children in the age range of this study (Diamond et al., 2007; Diamond & Kirkham, 2005).

### Working memory

Two subtests of the computerized Working Memory Test Battery for Children Aged Five to Twelve Years (AGTB 5–12; (Hasselhorn, 2012)) were used for the assessment of working memory. The AGTB 5–12 is a German standardized tool assessing working memory according to the multicomponent model by Baddeley (1986). Construct validity of the AGTB 5–12 was confirmed in a large study with 1,669 children (Michalczyk, Malstädt, Worgt, Könen, & Hasselhorn, 2013). We administered two subtests that are span measures with an adaptive testing procedure. We used the average score from the two subtests as a measure of working memory ability for each child. The subtests consist each of ten trials, each of them in turn divided into five testing blocks of two trials. The first testing block starts with a two-item sequence and sequence length is adjusted after each response: If the child recalls the presented trial correctly, the sequence length of the subsequent trial increases by one item. Following incorrect recall, the sequence length of the next trial is decreased by one item. In the remaining four testing blocks, the sequence length is adjusted more conservatively: If the child recalls both trials of a testing block correctly, the span length of the next block increases by one item. If, however, the child recalls both trials incorrectly, the span length decreases by one item (the minimum span length consists of two items). If recall is incorrect for only one of the two trials, the span length remains the same. The calculation of the span score is based on the mean performance in the last four testing blocks. For each correct response, the child receives a score that corresponds to the span length (i.e., the number of items within the presented sequence). For instance, if the child correctly recalls a five-item sequence, she receives five points. A false response is assigned the span length minus one (e.g., incorrect repetition of a five-item sequence results in four points).

Corsi span: Using a sequential presentation format, the Corsi span task captures the inner scribe of the sketchpad (e.g. Logie, 1995). In this task, a smiley face is displayed for 950 ms in one of nine white squares, placed pseudo-randomly on a grey background. After an inter-stimulus interval of 50 ms the smiley appears in a different square. At the end of each trial, children touch the squares where the smiley was shown, in the same sequential order.

Colour span backwards: The colour span backwards task captures two aspects of the central executive, namely coordinative complexity of controlling encoding and recall and selective focus on relevant information. A sequence of filled coloured circles is presented in the centre of the screen for 2 s each. At recall, children reproduce the sequence in reversed order by touching filled coloured circles (red, green, yellow, pink, blue, orange, black, brown) arranged in a larger circle.

### Reasoning ability

We operationalized reasoning ability using the Culture Fair IQ Test (Cattell, 1950). Briefly, the test consists of ten questions with where children see a series of images that follow a logical pattern. Because the pattern is never explicitly given, children have to infer it and choose the correct answer (out of five available) that is consistent with the inferred pattern. We considered the number of correct responses to the task.

### Behavioural data analyses

We did all analyses of behavioural data using R (v3.4.1, R Core Team, 2014), where we imported data initially stored in SPSS (package *foreign* (R Core Team et al., 2017); reorganized them (packages *plyr* (Wickham, 2016), *reshape2* (Wickham, 2017) and *stringr* (Wickham & RStudio, 2018)), built linear mixed models (package *lme4* (Bates et al., 2017)) and plotted them (package *ggplot2* (Wickham, Chang, & RStudio, 2016).

### MRI data acquisition

Structural data were acquired using a T1-weighted 3-D magnetization-prepared rapid gradient-echo sequence (repetition time = 2500 ms, echo time = 2500 ms, sagittal slice orientation, spatial resolution = 1 × 1 × 1 mm). T2*-weighted echo-planar images were acquired using a 3-T Siemens TIM Trio MRI scanner with a 12-channel head coil (transverse slice orientation, interleaved ascending scanning direction), field of view = 216 mm, repetition time = 2000 ms, echo time = 30 ms, 36 slices, slice thickness = 3 mm, matrix = 72 × 72, voxel size = 3 × 3 × 3 mm, distance factor = 10%, 152 volumes).

### MRI data analysis

### Preprocessing MRI data with the computational anatomy toolbox (CAT12)

We estimated cortical thickness using surface-based morphometry as implemented in CAT12 (r1278) (Jena University Hospital, Departments of Psychiatry and Neurology, http://www.neuro.uni-jena.de/cat/) running on SPM12 (Wellcome Department for Imaging Neuroscience, London, United Kingdom; http://www.fil.ion.ucl.ac.uk/spm) and Matlab R2016b (The MathWorks, MA, USA). We used an age-adequate tissue probability map (TPM), generated though the average approach of the Template-o-matic toolbox (Wilke, Holland, Altaye, & Gaser, 2008) instead of the default TPM for the segmentation, and default parameters otherwise. The data were then affine-registered to the MNI space and a non-linear deformation was applied. The deformation parameters were calculated with classical registration to the existing DARTEL template in MNI space generated from 555 participants from the IXI Dataset (Ashburner, 2007) (http://brain-development.org/ixi-dataset/). We did not correct the data manually, and a check of sample homogeneity revealed no issues. Surface and thickness were then estimated using projection-based thickness estimation methods (Dahnke, Yotter, & Gaser, 2013). Finally, we applied a smoothing kernel of 15 FWHM and submitted the resulting images to statistical analyses.

### Functional connectivity analyses

#### Preprocessing

We excluded the first five MR images of the series from the functional analyses to ensure steady-state longitudinal magnetization. We used SPM12 to preprocess the remaining images. We first performed slice timing correction and realignment, followed by coregistration between functional images and the individual anatomical T1 images. We then segmented the anatomical images into white matter, gray matter, and cerebrospinal fluid using the same age-adequate TPM as for the structural analyses. We normalized the resulting functional images to the MNI template and applied spatial smoothing with a 6-mm FWHM to improve signal-to-noise ratio. To reduce physiological high-frequency respiratory and cardiac noise and low-frequency drift, we used the REST toolbox (Song et al., 2011) to bandpass-filter (0.01 – 1 Hz) and detrend the data. We regressed out the signal from white matter and cerebrospinal fluid as well as the motion parameters. Additionally, we calculated the frame-wise displacement (FD) according to Power et al. (Power, Barnes, Snyder, Schlaggar, & Petersen, 2012). We excluded from the analyses one child who had an FD above the recommended threshold of 0.6.

### Functional connectivity and seed-based functional connectivity

To examine connectivity between brain regions using seed-based functional connectivity (FC) as implemented in the REST toolbox, we calculated the voxel-wise temporal (Pearson) correlations between a seed based on the structural results (see below) and the whole brain. We then applied Fischer transformations to the individual FC maps, to obtain z-scores to improve normality; and then submitted the z-score maps to a second-level analysis in SPM12, using movement (FD), age, sex, working memory, cognitive control and reasoning ability as covariates. We identified regions showing consistent levels of FC using a one-sample *t*-test (FWE corrected) with an additional threshold of *p*<0.05 at the voxel level, and cluster size >100 voxels.

## Results

### Behavioural results

Figure 2.A shows all responses to the knowledge and confirmation questions for each child and each condition. As Rohwer et al. (2012) reported, the *complete-knowledge* condition posed no challenge for children. Here, all 24 children answered (correctly) in both repetitions of the task that they knew which toy was in the box, but one child answered incorrectly in the confirmation question (i.e., they responded that they guessed the contents of the box, although they had seen the experimenter put the toy in the box). Although the *no-knowledge* condition was slightly more difficult, most (18 out of 24) children replied correctly to both instances. However, six children incorrectly responded in at least one instance that they knew which of two toys was inside the box, although they had not seen either of the toys. Finally, in the crucial *partial-knowledge* condition, 15 children responded (incorrectly) in at least one instance that they knew which of the two toys was in the box (and 9 children responded correctly to both instances). The *partial-knowledge* condition appeared to be the most difficult for children and in all but one case those children who responded incorrectly to at least one instance of the *no-knowledge* condition also responded incorrectly to the *partial-knowledge* condition.

If children answered (incorrectly) that they knew which toy was in the box in the *partial-* or *no-knowledge* conditions, the experimenter also asked how they knew this. Children often did not respond and, when they did, the answers included: “I don’t know”, “Because I’m smart” and “I know because I can read and hear”.

We studied the effect of age on performance in the partial knowledge task (figure 2.B) using a logistic regression. We found no significant effect of age (χ^2^(1)= 0.529, *p* = 0.467. In fact, the evidence favoured the null hypothesis of no effect of age, as estimated by a Bayes Factor with a standard Cauchy prior (BF*10* = 0.267).

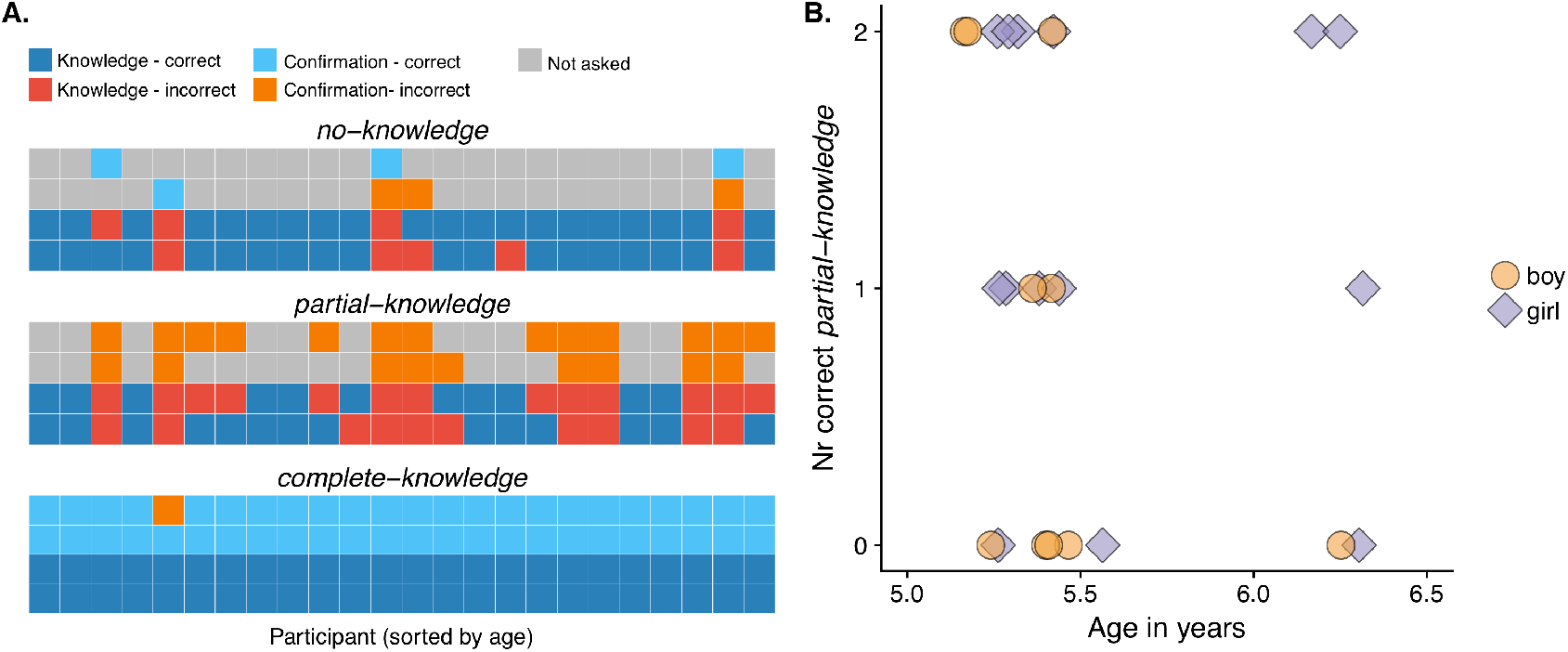

### Structural correlates of explicit metacognition

To identify brain structures that support meta-ignorance, we compared cortical thickness in the whole brain between two groups of children: those who responded correctly to both trials of the partial-knowledge condition and those who did not (similar to Wimmer & Perner, 1983). In a multiple regression, we looked for differences between the groups while accounting for age (in months), sex, working memory, cognitive control, and reasoning scores as covariates of no interest. After correction for non-stationary smoothness and expected cluster size, the group comparison revealed a positive effect in a single cluster spanning medial (79%) and superior orbitofrontal cortex (21%) (p = 0.00021, cluster extent k = 61, see figure 3.A). Further, two small but bilateral clusters showed a negative effect on superior parietal cortex (left: *p* = 0.00011, k = 30, 87% superior parietal, 13% inferior parietal; right: *p* = 0.00028, k = 15, 100% superior parietal, see figure 3.B).

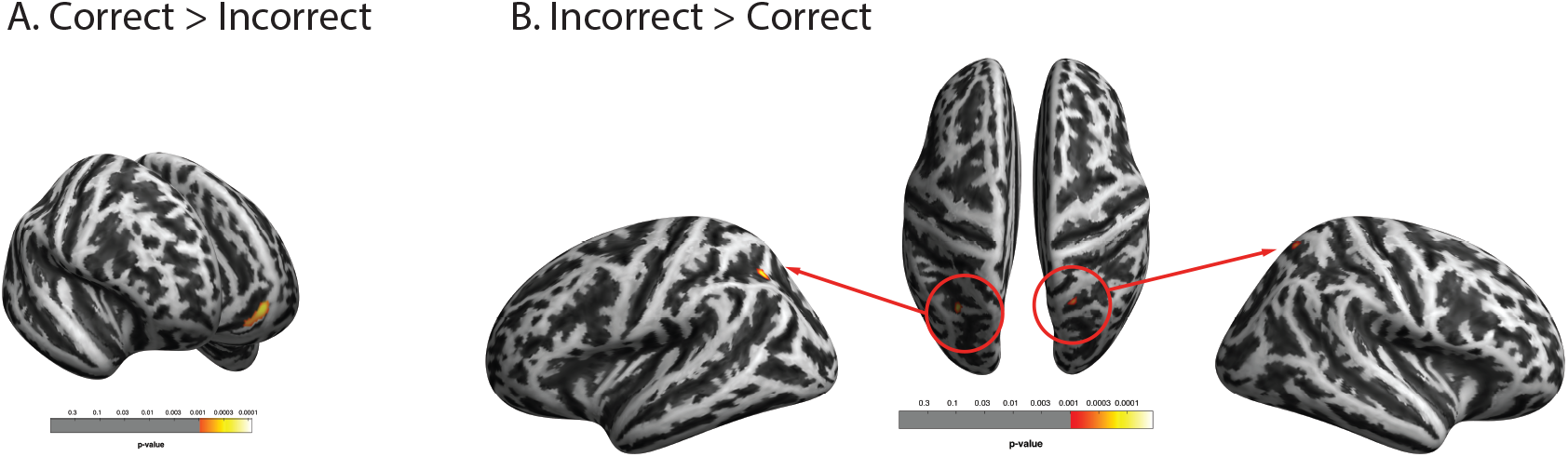

## Seed-based functional connectivity results

Structural analyses revealed that a region in the left medial orbitofrontal region is involved in the accurate processing of meta-ignorance. To better understand the role of this region, we aimed to identify the network that it is part of. We therefore measured seed-based connectivity during resting state, which infers the functional network between a region of interested given by a specific seed and all voxels in the brain. We calculated whole brain functional connectivity z-maps (based on Pearson’s correlations) for each child using a 10 mm-radius seed centred on (*x* = −8, *y* =, *z* = −1) and regressing out FD as a measure of movement (Power et al., 2012). We did a one-sample t-test and, as in the analysis of brain structure, we used age (in months), sex, working memory, cognitive control, and reasoning ability scores as covariates of no interest in the statistical model. Our group analysis showed statistically significant (i.e., consistent) functional connectivity between the prefrontal seed and left medial orbitofrontal gyrus, left posterior cingulate gyrus and right precuneus, and mid- and inferior temporal gyri (see table 1 and figure 4).

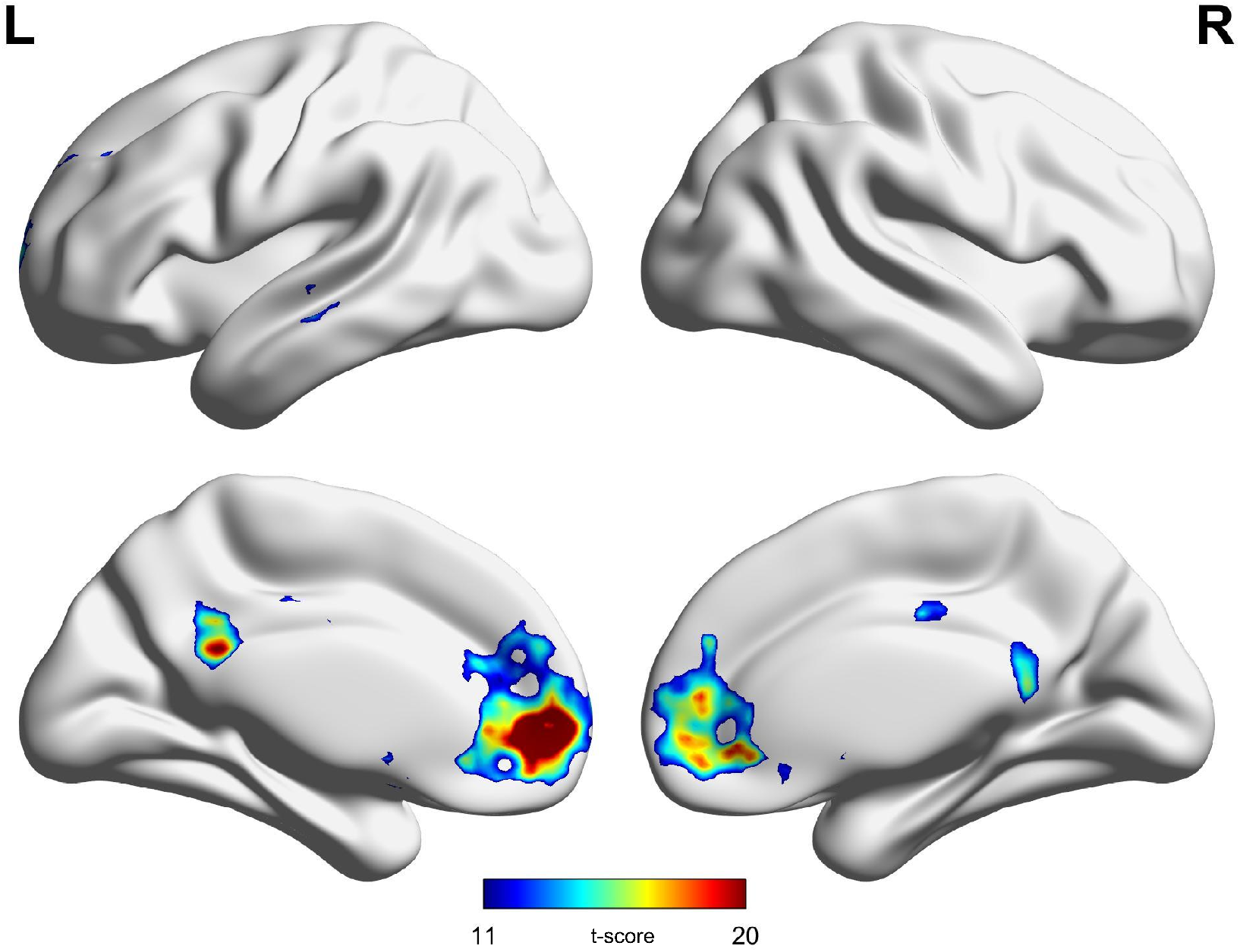

**Table 1:** Seed-based whole brain voxel functional connectivity results for the whole sample.

|  | Peak MNI Coordinates |  |  | *t*-score | Cluster size (voxels) |
| --- | --- | --- | --- | --- | --- |
|  | x | y | z |  |  |
| Left medial orbitofrontal cortex | -2 | 52 | -6 | 30 | 4676 |
| Left anterior cingulate gyrus | -12 | 48 | 0 | 30 |  |
| Left medial frontal gyrus | -10 | 58 | 2 | 28 |  |
| Left posterior cingulate gyrus | -4 | -42 | 24 | 21 | 919 |
|  | -10 | -46 | 30 | 17 |  |
| Right precuneus | 4 | -48 | 14 | 16 |  |
| Left middle temporal gyrus | -60 | -20 | -6 | 17 | 189 |
|  | -56 | -28 | -8 | 13 |  |
| Left inferior temporal gyrus | -52 | -22 | -20 | 13 |  |

## Discussion

We investigated the neural mechanisms relating to the emergence of explicit metacognition in preschool children by means of MRI. We concentrated on a “meta-ignorance” task, where children are required to recognize —and explicitly report— that they lack certain information. Previous results had shown that children develop this ability at around 5-6 years of age (Kloo et al., 2017; Rohwer et al., 2012). Our structural analyses revealed that a cortical region within the medial prefrontal cortex was thicker in those children that correctly identified that they did not know something. Our functional resting state analyses, in turn, revealed that this region is connected to the left medial orbitofrontal gyrus, left posterior cingulate gyrus and right precuneus, and mid- and inferior temporal gyri, which are regions belonging to the default-mode network (Deco, Jirsa, & McIntosh, 2011). These results highlight the neural networks that give rise to the emergence of explicit metacognition in early childhood. In the following sections, we bring our results in the context of the existing literature in both children and adults.

### Behavioural results

In their original study, Rohwer et al. (2012) used a cross-sectional design to test 2 to 7 year-old children’s performance in the meta-ignorance task. They found that the strongest differences in performance in the *partial-knowledge* condition occurred between 5 and 6 years-old children: there were no differences in performance between children of 2 to 5 years old, but children of all these ages performed worse than children aged 6 and 7. In our sample, we found that under half (9 out of 24) of the children aged 5-6 responded correctly to both repetitions in the *partial-knowledge* condition. We did not find a positive relationship between performance and age, but it should be noted that our age range was restricted. Overall, this corresponds to the findings by Rohwer et al. (2012) and demonstrates that we successfully implemented the task. This allows us to interpret the results of our structural and connectivity analyses.

### Structural results

We considered that a child is proficient in the meta-ignorance task if she could answer correctly to both instances of the *partial-knowledge* condition (similar to Wimmer & Perner, 1983). Nine out of 24 children could answer correctly to the two instances of this condition. We exploited this fact to build two groups: those that answered correctly to the meta-ignorance task and those who did not. We then compared cortical thickness estimates between these two groups, while accounting for other higher-order cognitive abilities like working memory, reasoning and cognitive control, as well as sex and age. We found significantly greater estimates of cortical thickness within left orbitofrontal cortex in the group of children that correctly answered to the meta-ignorance task. We also found that two small but bilateral clusters in the superior parietal cortex where cortical thickness estimates were significantly smaller in those children who responded correctly to the task. Both frontal and parietal regions have been described previously as relevant for performance monitoring and the formation of confidence (for a review, see Chua et al., 2014). Our study adds thus to previous research by demonstrating that these regions also subserve the emergence of metacognition. Here we discuss the two regions separately.

The effect we found in orbitofrontal cortex fits with the observation that structural parameters within the prefrontal cortex are associated with metacognitive ability (Bertrand et al., 2018; Cul, Dehaene, Reyes, Bravo, & Slachevsky, 2009; Filevich, Dresler, Brick, & Kühn, 2015; Fleming et al., 2010; McCurdy et al., 2013; Schnyer et al., 2004). Moreover, the direction of the effect (namely thicker cortex associated with better cognitive performance) is in line with neurodevelopmental trajectory studies showing that thickness increases in children with the age of our sample and peaks only later —at around 8 years of age— over the whole cortex (Raznahan et al., 2011) and in frontal regions specifically (Shaw et al., 2008). In particular, a recent study (Fandakova et al., 2017) on meta-memory found that cortical thickening in the ventromedial PFC predicted metacognitive improvements in 7 to 12 years-old children. Interestingly, our result also relates to research that addresses a very specific aspect of metacognitive monitoring: the ability to recognize that a stimulus has *not* been presented before. Miyamoto et al (Miyamoto, Setsuie, Osada, & Miyashita, 2018) recently showed that the bilateral frontopolar cortex (area 10) in macaques was causally related to metacognitive ability for correct rejections and false alarms (i.e., “noise trials”) but not to detection performance in the same trials or metacognitive ability for hits and misses (i.e., “signal trials”). Thus, our structural result, while being consistent with previous studies of different metacognitive abilities in adults, for the first time suggests a potential distinction in humans, akin to the distinction between detection and discrimination (Macmillan & Creelman, 2004) between the ability to monitor *that* one possesses a certain piece of evidence at all, and *which* information this is.

To the best of our knowledge, we are the first to report a negative relationship between metacognitive ability and structural cortical characteristics in the superior parietal cortex. The contribution of neural activity in primate parietal regions to decision confidence is well established (e.g., Kiani & Shadlen, 2009). In humans, BOLD signal levels in parietal cortex also vary with reported confidence, In particular, Chua et al (2014) have recently noted that, whereas inferior parietal cortex is often associated with high vs. low confidence, a handful of studies (Hayes, Buchler, Stokes, Kragel, & Cabeza, 2011; H. Kim & Cabeza, 2007, 2009; Moritz, Gläscher, Sommer, Büchel, & Braus, 2006) have reported that superior parietal cortex shows the inverse effect, i.e. higher BOLD signal levels in low confidence trials. On the basis of these results, Chua et al. hypothesised distinct roles of superior and inferior parietal cortex in confidence formation. Hence, while a link between functional results and structural characteristics is equivocal, we note that the negative relationship that we found between superior parietal cortex and meta-ignorance ability supports the hypothesis put forward by Chua et al. (2014). Future research may clarify the role of these regions in metacognitive ability.

### How does metacognition develop?

The behavioural paradigm to test meta-ignorance builds on an established line of research that examined meta-ignorance with respect to others’ or own knowledge (Rohwer, Kloo, & Perner, 2012; Sodian & Wimmer, 1987). Given the ongoing debate on the empirical relations between ToM and metacognition in preschool children (e.g., Lecce et al., 2015; Lockl & Schneider, 2007) and the theoretical dispute on the functional connection between ToM and metacognition (e.g., Apperly, 2008; Carruthers, 2009; Rohwer et al., 2012) we rely on existent activation-likelihood estimation (ALE) meta-analyses to compare the brain regions that we identified with the literature on the neural basis of ToM. Meta-analyses from studies in healthy young adults reveal ToM activations consistent across different tasks in the medial PFC, but these are mostly located ventrally to the cluster we identified (Schurz et al., 2014 -c.f. figure 5-; van Veluw & Chance, 2014). A meta-analysis that grouped together perceptual and memory metacognition identified several domain-general prefrontal regions: the posterior medial, dorsolateral, right anterior lateral and ventromedial PFC consistently showed higher levels of BOLD signal when collapsing across task modalities (Vaccaro and Fleming, *personal communication*). Hence, because —as far as these results show— the brain regions that support metacognitive monitoring differ from those supporting ToM, we suggest that these are distinct functions and that metacognition does not rely (solely) on the same neural architecture than ToM. Instead, these results favour the hypothesis that metacognition is related to networks supporting executive function (Roebers, 2017).

In order to better understand the role of the prefrontal region that we identified through structural analyses, we ran exploratory analyses to examine its functional connectivity pattern, which we describe below.

### Functional connectivity results

We used the cluster identified through structural analyses to calculate seed-based FC to explore consistent intrinsic functional connectivity patterns from the seed to the rest of the brain. We showed that the seed in left medial orbitofrontal cortex is functionally connected to left medial orbitofrontal gyrus, left posterior cingulate gyrus and precuneus, and mid- and inferior temporal gyri. From a network information-theoretic point of view, this result can be interpreted as a synchrony of the low-frequency components of the BOLD signal. Remarkably, the information processing from/to the medial orbitofrontal seed was seen in both long-range connections and local-range ones (neighbouring regions), which might integrate local information processing centres across the whole brain. This connectivity pattern includes three core regions of the default model network (DMN), namely the medial PFC (mPFC), posterior cingulate cortex/precuneus and lateral temporal cortices (Raichle, 2015). BOLD activity within DMN regions is typically associated with self-referential thought and introspection (Davey, Pujol, & Harrison, 2016; Northoff et al., 2006; Qin & Northoff, 2011). But, to the best of our knowledge, only a handful of studies have related functional connectivity during resting state to inter-individual differences in metacognitive ability before, though none of them has done so in a developmental population. First, Barttfeld et al. (2013) found that metacognitive accuracy in a visual task correlated positively with functional connectivity during an introspection condition correlated between the fronto-parietal system to itself as well as the sensorimotor system, the default brain mode network, and the cingulo-opercular network. Then, Soto et al (Soto, Theodoraki, & Paz-Alonso, 2017) found that functional connectivity between DMN regions and visual, right frontoparietal, and dorsal attention networks increased when participants introspected and recalled mental states during a cued visual search task that they had just done. And finally, Francis et al (2017) found that early-phase psychosis patients’ MAS-A scores (Metacognition Assessment Scale-Abbreviated, Lysaker et al., 2005) — which measures study several aspects of patients’ monitoring of self and others— correlated positively with functional connectivity between mPFC seeds and the posterior cingulate cortex and precuneus. Notably, despite strong differences in experimental design, all these studies point to two key common notions: They all identified the PFC as a key component of metacognitive monitoring, together with its connections to regions within the default mode network (DMN).

Notably, although each of these studies operationalized metacognition differently, they converge in reporting some involvement of the PFC in metacognitive monitoring. We caution that this should not be interpreted as the single atomic function of “metacognition”, nor that PFC is a functionally homogeneous brain region. In fact, Baird et al (2013) have shown that that two seed ROIs defined *a priori* within the anterior PFC had different connectivity patterns and, crucially, correlated differently with ability in two separate metacognitive tasks. In light of this discussion, one important question is whether, and to what extent, the DMN in our developmental sample is analogous to that of adults. Overall, the existing literature indicates that an adult-like DMN architecture is already present and relatively stable by around 1 year of age. In particular, the posterior cingulate and retrosplenial cortices, as well as and mPFC appear to be crucial network hubs and are integrated into it already in the newborn brain (Gao et al., 2009). More specifically, at least two studies measured functional connectivity during resting state in infants (41 weeks-old babies born at extremely low gestational age (Fransson et al., 2007) and 2-4 weeks-old newborns (Gao et al., 2009)). Both studies found that the brain regions active during rest in infants overlap only partially with the adult DMN. However, functional connections (in particular long-distance) develop rapidly during the first year of age (Gao et al., 2009), and continues to develop at a slower pace for several years after that (Gao et al., 2011) Fair et al. (2008) studied 7-9 years-old children and adults and found that, although the mPFC was part of the DMN in all age groups, the connections between mPFC and posterior cingulate and parietal regions were weaker in children. This was not only the case for these connections. In fact, although the network consisted of the same regions, these were sparsely or weakly connected in children as compared to adults. Further, Khan et al (2018) reported that the DMN continues to mature between at least 8 years old and adulthood on a frequency-band-dependent fashion. Different sets of regions showed changes in different frequency bands, and they showed different temporal patterns. In short, the DMN network continues to mature and develop into adolescence, so we caution against committing to an unqualified interpretation of these results. But, to the extent that the main features of the network are already in place at the age of the children in our sample, we are justified in interpreting the functional connectivity results in light of self-reflection and DMN. Interestingly, our results contribute to the body of literature linking metacognition to self-referential thought, thus suggesting a potential mechanism by which the brain monitors itself. To further test this notion, future studies could investigate functional connectivity during rest vs. a cognitive task in the same children that do a metacognitive task.

### Limitations

Two main limitations of these results should be mentioned. First, as virtually any empirical study of metacognitive processing, the results cannot *a priori* be generalized to other metacognitive functions. Just as studies in adults have revealed different neural bases for different metacognitive tasks and suggested some degree of domain-specificity (Baird, Mrazek, Phillips, & Schooler, 2014; Fleming, Ryu, Golfinos, & Blackmon, 2014), studies in developmental populations have revealed that different aspects of metacognitive monitoring develop at different developmental stages, namely implicit and explicit monitoring, (Goupil & Kouider, 2016; Goupil et al., 2016; S. Kim et al., 2016). We note that inter-individual differences between these different aspects may still be stable over development. To the best of our knowledge, this has not been directly investigated and could be the focus of future studies.

With this in mind, we now consider three potential specific objections to the experimental paradigm. First, could these results be explained by the parsimonious interpretation that children had a general bias against admitting their own ignorance, or that they did not understand the terms “knowing” and “guessing”? The results of the *no-knowledge* condition argue against this explanation: If any of these two scenarios were true, those children that answered incorrectly in the *partial-knowledge* condition should have also answered incorrectly in the *no-knowledge* condition (i.e. answering that they knew which toy was in the box in both cases). This was only the case for five out of the 15 children in our sample, so this effect alone cannot explain our results. Further, while we did not evaluate whether the children in our sample understood these concepts well, the existing literature suggests that children this age understand the difference between (and spontaneously use) the terms “knowing” and “guessing” without ambiguity (e.g., Moore, Bryant, & Furrow, 1989; Shatz, Wellman, & Silber, 1983). See Rohwer et al. (2012) for a more detailed argument.

Second, our analysis of children’s behaviour in the meta-ignorance task followed Wimmer and Perner (1983) by treating metacognitive ability as binary: a child can either solve the problem, providing consistently correct answers, or she cannot yet. This methodological step allowed us to maximize the contrast and explore the neural bases of metacognition. Yet, metacognitive ability is not all-or-none and its development does not finish during childhood. Other paradigms that quantify metacognitive ability not just as a categorical variable (correct or incorrect, as in this study) but rather as a continuous measure (i.e., the accuracy of confidence ratings following correct responses) may offer finer grained information (Geurten, Willems, & Meulemans, 2015; Hembacher & Ghetti, 2014; Paulus, Tsalas, Proust, & Sodian, 2014; Weil et al., 2013; see also Ghetti, Hembacher, & Coughlin (2013) for a review). Future studies may consider these paradigms in early developmental samples like ours to draw connections between the developmental trajectory or metacognitive ability and its neural bases in the normal adult brain.

A final limitation of these results is our small sample size. We nevertheless note that the imaging results that we report here are consistent internally and with the existing literature. As we argued above, the prefrontal cluster where cortical thickness correlated positively with meta-ignorance ability matches those previously found in the literature in relationship with metacognition, and in particular the monitoring of non-experienced stimuli. Additionally, the cortical thickness results were consistent with those from the functional connectivity data during resting state. All in all, our results are consistent and go beyond previous literature, and may help constrain and inform future studies.

### Conclusion

We used a meta-ignorance task to identify the structural and functional correlates of metacognitive ability in children at the age where this ability develops. Children who answered correctly to a metacognitive task had greater cortical thickness in a cluster within medial prefrontal cortex, compared to children who answered incorrectly to the task. Functional connectivity analyses revealed that this region connects to other frontal regions, medial orbitofrontal gyrus, posterior cingulate gyrus and precuneus, and mid- and inferior temporal gyri. The cluster within medial prefrontal cortex did not overlap with clusters previously identified in adults as supporting theory of mind, a cognitive function thought to be related to —and, under some accounts, be a precursor of— metacognition. Instead, the complete pattern of results recalls the default-mode network, often associated with self-referential thought and introspection. Our results suggest that children’s metacognitive ability to recognize that they do not know something depends on a mature default-mode network that supports introspective processing.

## Acknowledgements

We thank Yana Fandakova for very helpful comments on an earlier version of this manuscript. This work was supported by a Minerva Research Group to YLS from the Max Planck Society. YLS has been funded by the European Union (ERC-2018-StG-PIVOTAL-758898) and a Fellowship from the Jacobs Foundation (JRF 2018–2020). EF and CF are supported by the Volkswagen Foundation (grant number 91620). SK has been funded by two grants from the German Science Foundation (DFG KU 3322/1-1, SFB 936/C7), the European Union (ERC-2016-StG-Self-Control-677804) and a Fellowship from the Jacobs Foundation (JRF 2016-2018). MP has been supported by a Fellowship from the Jacobs Foundation (JRF 2016 1217 12). The design of the study, the collection, analysis, and interpretation of the data, and the writing of the manuscript was the sole responsibility of the authors.

